# Oscillatory markers of interoceptive attention: beta suppression as a neural signature of heartbeat processing

**DOI:** 10.1101/2025.06.24.660939

**Authors:** Kristina Pultsina, Suvi Karjalainen, Tiina Parviainen

## Abstract

Interoceptive attention—the ability to selectively focus on internal bodily signals—has been linked to distinct neural responses, yet the contribution of oscillatory dynamics to this process remains underexplored. This study investigates the neural mechanisms underlying interoceptive attention by examining beta-band power suppression during heartbeat and auditory discrimination tasks. Fifty-one healthy participants engaged in interoceptive (heartbeat detection) and exteroceptive (auditory discrimination) tasks while their brain activity was measured using magnetoencephalography (MEG). The results revealed significant beta suppression time-locked to the R-peak in the somatosensory cortex, anterior cingulate cortex, mid-cingulate cortex, and dorsolateral prefrontal cortex from 310 to 530 ms post-R-peak. Beta suppression was more pronounced during interoceptive attention, correlating positively with interoceptive accuracy. The findings support the notion that beta suppression in fronto-cingulo-somatosensory network may serve as a neural marker of interoceptive processing, contributing to predictive coding models of interoception. This study highlights the potential for using beta suppression as an objective measure of interoceptive accuracy and suggests that neural oscillations play a critical role in the brain’s regulation of heartbeat-related information. Furthermore, the study proposes that interoceptive attention involves a top-down mechanism that dynamically adjusts the brain’s response to cardiac afferent signals, enhancing the precision of interoceptive processing. These findings have implications for understanding how the brain integrates interoceptive signals and may provide insights into clinical applications targeting interoceptive dysfunctions.

## Introduction

Interoception refers to the perception of sensory information originating from the viscera and internal milieu. The ability to attend to and consciously process internal bodily signals is considered crucial for fundamental aspects of selfhood and emotion regulation, as well as for optimal cognitive functioning from basic goal-directed behavior to higher-order executive tasks. A broader definition of interoception involves not only low-level processing of individual visceral signals, such as changes in the biochemical composition of blood, but also the integration of information about physiological state, which in turn can influence cognitive processes such as attention and memory, as well as emotions (Craig, 2009; Critchley & Harrison, 2013; Tsakiris & Critchley, 2016).

Cardiac interoceptive sensitivity, specifically the ability to discriminate one’s own heartbeats, serves as a paradigmatic example of this phenomenon. Elucidating the physiological pathways underlying cardiac interoceptive sensitivity is crucial for comprehending the complex interplay between cardiac and neural systems, and its role in individual differences in experience. A potential source of neural responses to heartbeats is the activation of mechanoreceptors in the heart wall, aortic arch, and carotid sinus, along with cutaneous somatosensory mapping and neurovascular coupling in the cortex (Park & Blanke, 2019; Tallon-Baudry et al., 2018). This information travels via the nucleus of the solitary tract and then mainly via thalamus, functionally connected to forebrain regions, including amygdala, insular cortex, anterior and mid cingulate, and orbitomedial prefrontal cortex (Critchley & Harrison, 2013).

Neuroimaging studies have implicated the insular cortex and anterior cingulate cortex (ACC) as pivotal structures in the representation of interoceptive states, including the modulation of cardiac activity (Craig, 2009). The ACC’s involvement in both central autonomic control and peripheral sympathetic responses across various contexts suggests its role in integrating visceral states with behavioral output (Craig, 2009; Critchley et al., 2011).While the insula is predominantly associated with generating emotional arousal and the cortical input of interoceptive signals into the brain, the anterior cingulate cortex, known as the ’limbic motor cortex’ is crucially involved in generating autonomic responses to emotionally salient stimuli (Craig, 2009; Medford & Critchley, 2010; Uddin, 2015). Notably, different studies have shown increased activity in these brain structures when participants were asked to attend to internal bodily responses, for example, to their heartbeats in Heartbeat Detection task (Critchley et al., 2004a; Petzschner et al., 2019; Pollatos et al., 2005).

Recent research suggests the insula and ACC are essential for interoceptive awareness, but a study by Khalsa challenges this view (Khalsa et al., 2009). Using isoproterenol to induce heartbeat sensations, they found that a patient with bilateral insula and ACC damage could still perceive his heartbeat, like healthy controls. However, when his skin sensations were blocked by a topical anesthetic, his interoceptive awareness disappeared. These results suggest that both visceral and somatosensory pathways contribute to interoception, rather than the insula alone. This is consistent with neuroimaging findings showing that activation of the somatosensory cortex correlates with interoceptive awareness in tasks requiring focused attention on the heartbeats (Critchley et al., 2004b; Pollatos et al., 2007).

The cumulative evidence supports the existence of multiple, semi-independent pathways subserving cardiac interoceptive awareness and suggests a complex interplay between somatosensory processing and interoceptive phenomena. This is consistent with the idea of neurovisceral integration that interoception is part of a broad multisensory integration used for self-regulation and adaptability of the organism (Pezzulo et al., 2015; Thayer & Lane, 2000). The precise neural mechanisms involved in integrating interoceptive signals remain largely unexplored. According to the influential predictive coding theory, top-down predictions about the body’s state are thought to be continuously compared with incoming bottom-up sensory information (Owens et al., 2018).

The most common marker of heart–brain communication is heart beat evoked potential (HEP), which reflects the brain’s processing of cardiac signals. Even though evoked potential predominantly reflects bottom-up processing of sensory signals, in this case, the brain processes heartbeat signals through baroreceptor activity (Park & Blanke, 2019; Suarez-Roca et al., 2021). There is also evidence that attention focus and affective predictions about external stimuli can influence HEP amplitude (Al et al., 2020; Gentsch et al., 2019; Petzschner et al., 2019).

There is growing evidence that bodily sensations have been linked to modulations of induced activity. For example, breathing cycles modulate alpha rhythms in the brain, both at rest and during both spontaneous and deep breathing (Karjalainen et al., 2025; Kluger et al., 2021; Zaccaro et al., 2025). Similarly, arousal has been shown to modulate the cortical processing of heartbeats, with higher arousal states amplifying HEPs. Notably, alpha oscillations influence individual variability in interoceptive sensitivity, suggesting a potential gating mechanism for cardiac awareness. This highlights the crucial role of oscillatory activity in shaping the brain’s dynamic integration of bodily signals (Luft & Bhattacharya, 2015).

Moreover, recent evidence highlights the role of cortical oscillatory dynamics in cardiac interoception. Zaccaro et al. demonstrated that heartbeat-related alpha and theta band oscillations, particularly during exhalation, are associated with increased interoceptive accuracy during heartbeat counting tasks, with these oscillations – reflected in power, inter-trial coherence, and functional connectivity – being most prominent over right fronto-centro-parietal regions and independent of peripheral cardiac physiology (Zaccaro et al., 2025). Importantly, alpha-band modulations showed a significant correlation with heartbeat-evoked potentials, suggesting that oscillatory activity may support the top-down attentional allocation to cardiac signals by enhancing the precision-weighting of prediction errors.

The study by Crivelli & Balconi found that interoceptive attention during the Heartbeat Counting Task (HCT) was associated with increased alpha oscillatory activity in the right parahippocampal gyrus, suggesting that alpha rhythms play a role in top-down attentional control, helping to suppress irrelevant external information while focusing on internal bodily signals (Crivelli, D., Balconi, 2025). Additionally, beta oscillations in the cingulate cortex were positively correlated with interoceptive accuracy, indicating that beta activity reflects active processing of interoceptive signals rather than just passive bodily awareness. Notably, these neural markers were not linked to heart rate variability, reinforcing the idea that interoceptive attention is guided by cognitive control rather than direct cardiac input. These findings support predictive coding models of interoception, highlighting oscillatory induced activity as a potential objective marker for interoceptive accuracy and regulation.

Thus, the analysis of induced oscillatory activity may provide a better understanding of the mechanisms of predictive processing, adaptation, and integration of interoceptive information over time. Specifically, we focused on oscillatory dynamics in the beta frequency bands, hypothesizing that directing attention to heartbeats would modulate these rhythms – in particular, that it would produce R-locked power changes as a neural signature of interoceptive attention in the heartbeat and auditory discrimination task. The primary focus is on beta-band power, given its critical role in the top-down control of predictions, with a decrease in beta power associated with updating predictions and processing prediction errors (Engel & Fries, 2010).

## Results

To assess the presence of R-peak-related changes in beta band, we performed a one-sample cluster t-test on all vertices for the collapsed exteroceptive and interoceptive tasks.

A significant TFCE cluster was identified between 310 ms and 530 ms post-R-peak with a pronounced suppression at beta frequency band in anterior (ACC) and middle cingulate cortex (MCC), somatosensory and premotor cortex (S & PMC), as well as the dorsolateral prefrontal cortex (dlPFC) (Fig.2).

**Fig.1.**
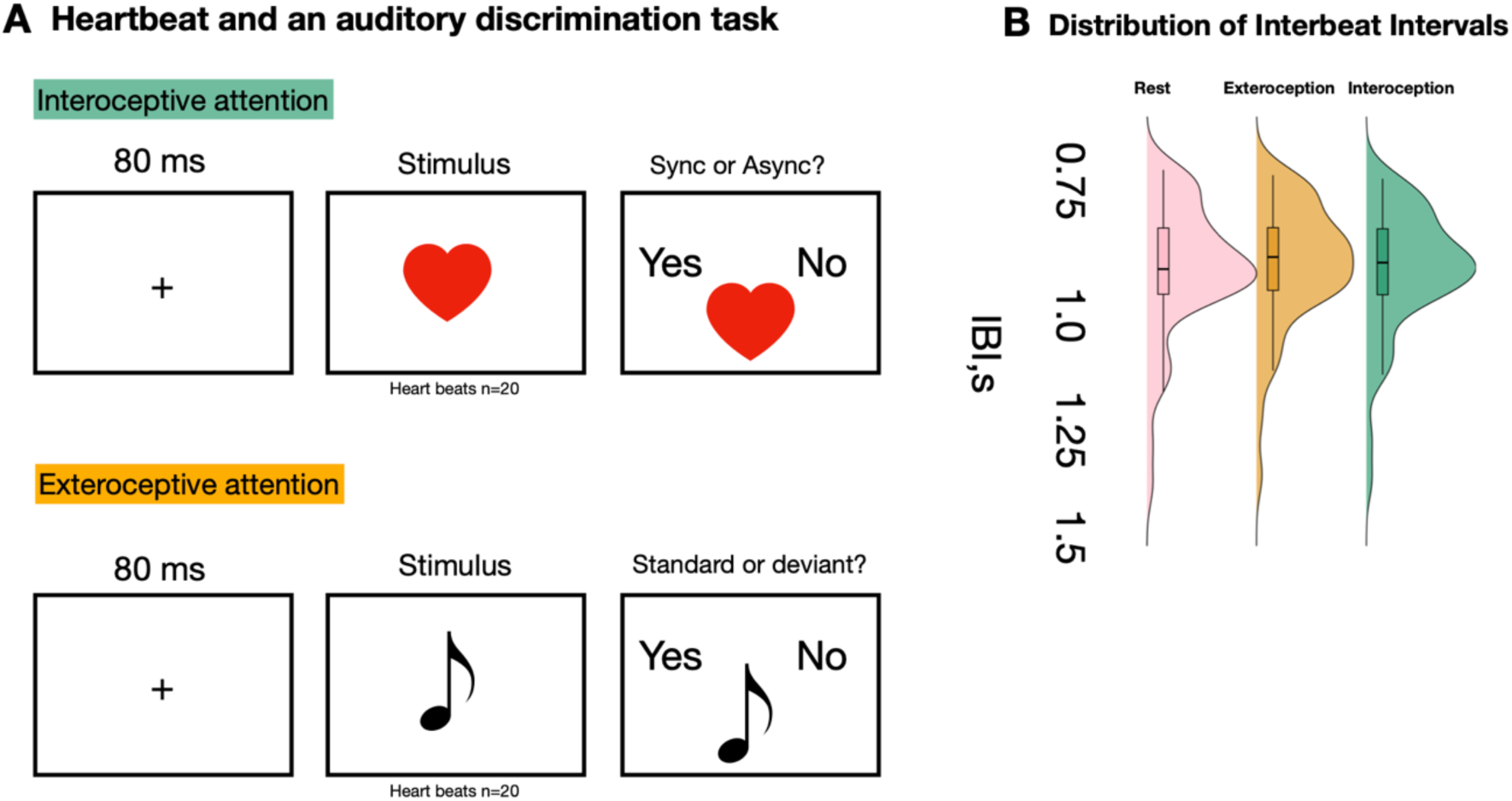
Schematic overview of the block design in the heartbeat detection and an auditory discrimination task. **A** The task alternated between two types of attention blocks: the interoceptive condition, where participants focused on their heartbeat, and the exteroceptive condition, where they attended to the sound presented through headphones. **B** Distribution of Interbeat Intervals (IBI) Across Rest, Exteroception, and Interoception Conditions.

**Fig.2.**
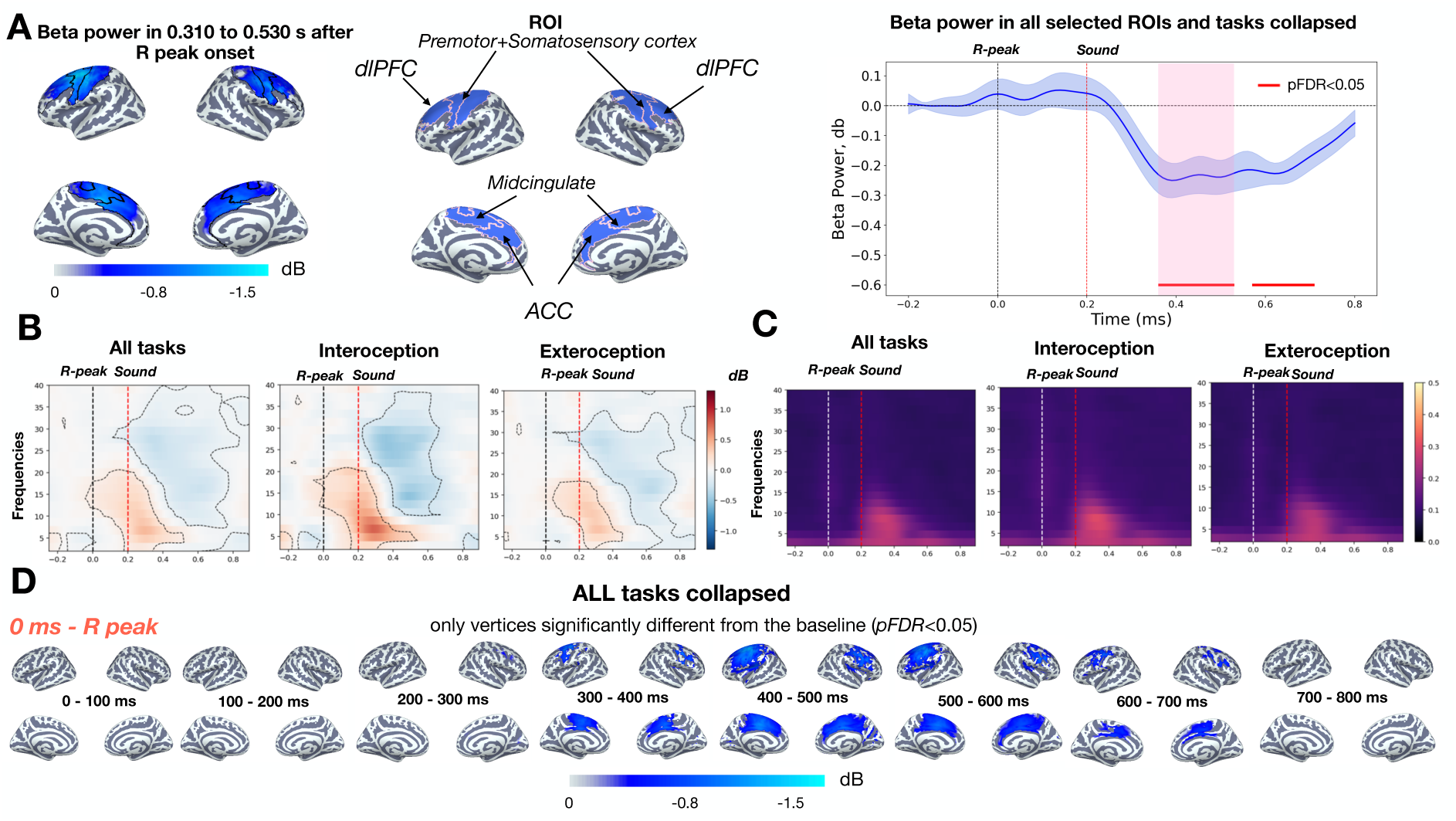
R-peak related β suppression in all conditions collapsed. **A** - Source distribution of β power in the TFCE defined cluster time interval 0.310 to 0.530 s (only significant vertices shown) for all tasks combined; ROIs with newly defined borders thresholded by significant vertices; Time course of β power changes across all tasks and ROIs. **B** - Time-frequency distribution across all pooled ROIs. The dashed line indicates the area that differs significantly from the baseline (one-sample t-test, pFDR<0.05). **C** - Phase-locked distribution across all pooled ROIs. **D** – Beta power changes following the R peak, averaging over 100-ms intervals. Only vertices that significantly differ from the baseline are shown (one-sample t-test, pFDR < 0.05).

Further refinement of the preselected ROI boundaries (see Methods) identified statistically significant vertices in all regions except for the insula and frontal operculum cortex, which were excluded from further analysis (Fig. 2A). The time-frequency analysis (Fig.2 B,C) showed that besides modulation at beta band, also alpha band (8 - 14 Hz) suppression was related to heartbeats. We performed the same analysis to check for significant alpha suppression but found none.

### Beta power during interoception and exteroception tasks

After identifying the main ROIs with significant suppression, we performed a comparison of oscillatory activity at beta band between the exteroceptive and interoceptive tasks using the average power in the time window of 0.310 to 0.530 s after the R-peak.

We found significant differences between the interoception and exteroception tasks for all the selected ROIs (MCC: t(50) = 3.967, p<0.001, d = 0.556; S & PMC: t(50) =2.933, p=0.005, d = 0.411; dlPFC: t(50) =3.872 p<0.001, d = 0.542; ACC: t(50) =3.857, p<0.001, d = 0.540). During interoception, participants demonstrated more prominent suppression in the somatosensory cortex and premotor cortex, dlPFC, ACC, and midcingulate cortex (Fig.3A). Also, to validate that the observed difference in suppression was not whole-brain related, we performed the same comparison between the tasks on the whole-brain averaged data and didn’t reveal any significant differences (t (50) =-0.517, p=0.608, d = 0.072).

**Fig.3.**
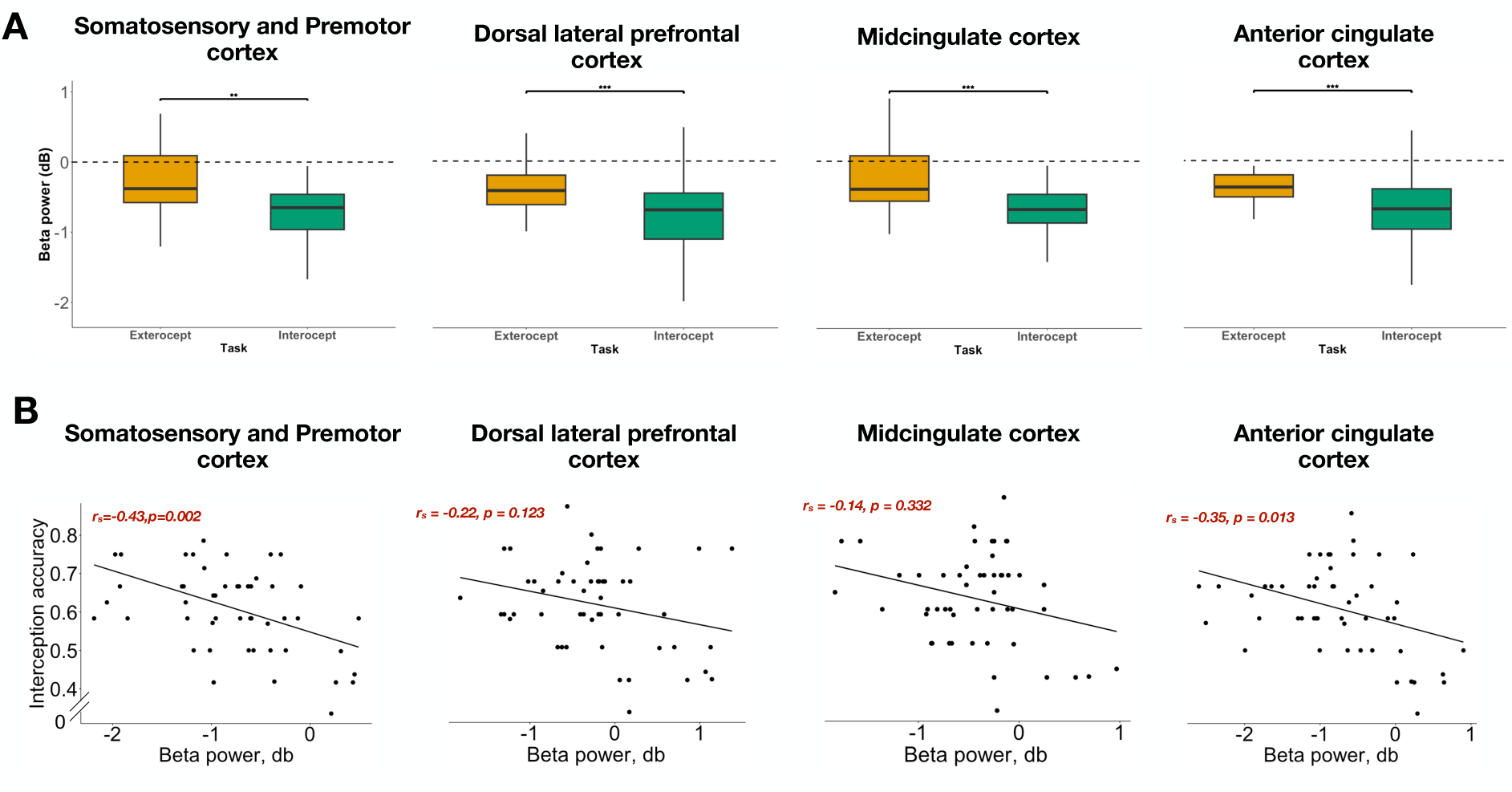
Beta power change in the ROIs averaged in 0.310 to 0.530 s after R-peak onset. **A** - Boxplots comparing mean beta power in exteroceptive and interoceptive attention tasks (**p ≤ 0.01, ***p ≤ 0.001). **B** - Correlation between beta power during interoceptive attention task and the interoception accuracy.

To rule out the possibility that the observed differences between tasks were due to difference in duration in heartbeat intervals, we compared the heart interbeat intervals (IBI) during interoception and exteroception task (Fig.1B). The t-test did not show significant difference between exteroception and interoception task (t (50) = - 0.458, p=0.649, d =0.064) in IBI. The average IBI for the exteroception task was 0.926 s ± 0.018 and 0.930 s ± 0.018 for the interoception task.

### Correlation between interoception accuracy and beta suppression during the interoception task

To test whether beta suppression in ROIs related to interoceptive accuracy, we performed a correlation analysis. A significant correlation between accuracy and beta power suppression was found for the S & PMC (r_s_ (49) = −0.43, p=0.002) and ACC (r_s_ (49) = −0.35, p = 0.013) ROIs. Higher accuracy was related to greater suppression in these brain areas. The same trend was observed in the anterior and mid-cingulate cortex; however, the correlation values did not reach statistical significance.

### Beta suppression during resting state condition

We further tested if the observed suppression is specifically related to the tasks requiring attention to exteroceptive or interoceptive signals, or whether the task instead modulates R-peak suppression that is already present at rest?

To investigate this, we used resting state data. We averaged beta power within the regions of interest (ROIs) and the time interval that exhibited significant suppression during both the interoceptive and exteroceptive tasks. We then compared these values to the baseline using a one-sample t-test.

The results revealed significant beta suppression in somatosensory and premotor ROI (t (50) = −3.058, p=0.004, d=0.428), MCC ROI (t (50) = −2.982, p=0.004, d=0.418) and dlPFC ROI (t (50) = −3.068, p=0.003, d=0.430). However, in the ACC, beta suppression did not reach statistical significance (t (50) = −1.947, p=0.057, d=0.273) (S1 Fig).

The comparison of resting-state beta suppression and during interoception and exteroception tasks was analyzed using a rmANOVA. The results showed a significant effect of condition on beta power in the S & PMC (F (2,50)=14.76, p<0.001, ηp2= 0.228), MCC (F (2,50) =21.39, p<0.001, ηp2= 0.3) and dlPFC (F (2,50)=25.18, p<0.001, ηp2 =0.335). Post-hoc comparisons revealed the lowest beta activation in the resting state (S & PMC: Rest vs Exteroception p=0.011, Rest vs Interoception p<0.001; MCC: Rest vs Exteroception p=0.021, Rest vs Interoception p<0.001; dlPFC Rest vs Exteroception p<0.001, Rest vs Interoception p<0.001) (S2 Fig.).

### Temporal Specificity of R-peak related beta suppression

To determine whether the observed post-R-peak beta-band suppression is time-locked to cardiac events – reflecting neural responses to afferent signals following heart contraction— rather than being related to motor activity or spontaneous neural fluctuations, we generated surrogate epochs for each participant (Park et al., 2014; Steinfath P , Herzog N , Fourcade A , Sander C , Nikulin V, 2024). This approach allowed us to test whether the beta suppression was specifically aligned with actual cardiac events rather than with efferent processes preceding the heartbeat or random fluctuations. However, when using the surrogate R-peak epochs, this post-R-peak beta suppression vanished in all ROIs, demonstrating that the effect requires precise alignment with true heartbeats (S3 Fig). In other words, randomly shifting the timing of heartbeats negated the beta suppression effect, yielding no significant post-trigger power drop in the beta band.

We further examined whether the timing or magnitude of post-R-peak beta suppression depended on the length of the cardiac cycle, which would be expected if the brain were anticipating the next heartbeat. Heartbeat events were divided into two groups based on the interbeat interval (short vs. long, via a median split of IBI durations). We then compared the post-R-peak beta activity between these groups. The results showed no significant difference in the onset, duration, or amplitude of beta suppression between short-IBI and long-IBI heartbeats – the suppression followed the preceding R-peak on a consistent timeline in both cases (S4 Fig). This temporal invariance directly contradicts an anticipatory explanation. If the beta suppression were an anticipatory response to the upcoming beat, we would expect it to adjust based on the expected timing of that beat (for example, occurring earlier or ending sooner when the next heartbeat is imminent). Instead, the beta suppression appears to be locked to the previous heartbeat’s afferent signals and does not shift in accordance with the varying interval to the next beat. Such consistency indicates that the suppression is driven by the physiological consequences of the preceding cardiac cycle (e.g. baroreceptor input from that heartbeat) rather than by a countdown or prediction of the next heartbeat’s timing.

## Discussion

Our study, using a heartbeat and an auditory discrimination task, provides evidence that paying attention to internal bodily signals – in this case, one’s heartbeat – engages distinct neural dynamics characterized by suppression of beta-band oscillatory activity in the premotor and somatosensory cortex, anterior and medial cingulate cortex, and dorsolateral prefrontal cortex. In other words, when participants focused on feeling their heartbeat, the beta rhythms (16 – 29 Hz) significantly decreased relative to when they attended to external sounds. Intriguingly, the depth of beta suppression positively correlated with interoceptive accuracy.

This finding supports existing evidence that the focus of attention modulates interoceptive signal processing (Critchley et al., 2004a; Petzschner et al., 2019). Compared to previous studies that primarily examined phase-locked heartbeat-evoked responses or focused on alpha- and theta-band modulations during interoceptive attention (Lee et al., 2024; Petzschner et al., 2019; Zaccaro et al., 2025), our findings extend this body of work by demonstrating that dynamically shifting attention focus -specifically when integrating internal cardiac signals with external auditory information – engages distinct oscillatory mechanisms, particularly beta-band suppression. This interpretation is consistent with previous evidence linking beta suppression with integrating sensory input with existing priors and the maintenance of ongoing cognitive processes (Arnal & Giraud, 2012; Kilavik et al., 2013; Sherman et al., 2016).

While several studies have reported alpha-band changes during interoceptive attention, particularly in heartbeat counting tasks that require sustained internal focus (Crivelli, D., Balconi, 2025; Villena-González et al., 2017; Zaccaro et al., 2025), our findings instead revealed prominent beta-band suppression. The observed differences in results can be attributed to several factors. Primarily, they stem from variations in the experimental task: participants engaged in a heartbeat counting task, which relies on internally directed, top-down attention and the inhibition of brain areas not directly involved in the task (Crivelli, D., Balconi, 2025; Zaccaro et al., 2025). This form of attention has been associated with increased alpha-band activity, reflecting the suppression of external input. The alpha synchronization observed in visual and parietal regions during interoceptive attention, relative to attention to external stimuli, likely reflects a neural gating mechanism that filters out task-irrelevant information (Pfurtscheller et al., 1996; Villena-González et al., 2017).

By contrast, our heartbeat and an auditory discrimination task required splitting attention between internal signals and external tones, reflect a shift away from a stable, “status quo” state toward dynamic processing (Engel & Fries, 2010). The R-peak locked beta suppression we observed reflects this active engagement, where each heartbeat-tone pair demands evaluation, eliciting transient beta desynchronization. Within the predictive coding framework, heartbeat-related beta suppression may reflect a comparison between priors about interoceptive signals and incoming sensations (sound). For instance, hearing one’s own heartbeat evokes a smaller auditory N1 response than hearing external sounds, reflecting predictive attenuation of expected input (Van Elk et al., 2014). In this context, beta suppression may serve as a neural mechanism for minimizing the impact of predictable interoceptive stimuli, consistent with classical sensory gating and predictive coding accounts. Notably, the fact that suppression was independent of cardiac cycle length further supports its top-down, cognitive nature, rather than a purely physiological reflex.

Finally, an important piece of evidence linking neural mechanisms to function is that the strength of heartbeat-related beta suppression in ACC, somatosensory, and premotor areas correlated with interoceptive accuracy. This supports the interpretation that beta suppression indexes interoceptive signal processing. Prior studies have similarly linked interoceptive accuracy to oscillatory brain activity (Pollatos & Schandry, 2004; Zaccaro et al., 2025). For example, prior studies have shown that individuals with higher heartbeat detection accuracy tend to exhibit larger HEP amplitudes, and that beta oscillation changes in the cingulate gyrus during heartbeat counting tasks are significantly associated with interoceptive performance (Crivelli, D., Balconi, 2025; Pollatos & Schandry, 2004). Together, these findings support the notion that beta suppression may serve as a candidate neural marker of interoceptive sensitivity, reflecting the intentional engagement of attention toward subtle internal sensations rather than basic, low-level sources of information, such as heartbeat-related sensations; however, this interpretation requires further validation.

The spatial distribution of beta suppression – especially in the anterior cingulate cortex, somatosensory cortex, and prefrontal areas – partially aligns with earlier fMRI data on brain structures engaged during interoceptive attention (Critchley et al., 2004a; Pollatos et al., 2007). One notable divergence from prior neuroimaging studies is the absence of significant heartbeat-related effects in the insular cortex. This may reflect the insula’s role in sustained or integrative interoceptive processes that are not captured by time-locked analyses. fMRI, for instance, can detect overall increases in insular activity during interoceptive tasks, but lacks the temporal resolution to resolve rapid, R-peak-locked oscillatory changes. It is conceivable that insular activation reflects a meta-representation of bodily state or subjective feeling of the heartbeat, rather than transient responses to individual cardiac afferent bursts (Corbetta et al., 1991; Craig, 2009). Second, technical factors could have limited our ability to detect insular signals. Additionally, the insula’s deep and folded anatomy may limit MEG/EEG sensitivity, further contributing to the absence of detectable effects (Hauk et al., 2022). Therefore, our findings do not preclude insular involvement but highlight the need for complementary imaging approaches to fully capture its contribution to interoception.

Importantly, the cortical sources we identified for heartbeat-related beta suppression are consistent with the proposed “second pathway” of cardiac interoception that involves somatosensory processing (Khalsa et al., 2009; Park & Blanke, 2019). Our results resonate with the case of Khalsa’s patient and other evidence suggesting that cardioceptive signals can be routed through somatosensory cortices (Khalsa et al., 2009). Observed activation in the somatosensory and motor cortex, MCC, and ACC may be related not only to the direct monitoring of heartbeats through baroreceptors – connected to the nucleus tractus solitarius (NTS) – but also to peripheral changes associated with heartbeats (Critchley & Harrison, 2013).

Thus, our findings reinforce the view that interoceptive awareness of heartbeats is the product of an integrated viscerosensory and somatosensory process.

Recent neurophysiological studies provide compelling evidence for a bidirectional interplay between central neural oscillations and peripheral autonomic activity, particularly involving the dlPFC, ACC and somatosensory cortex (James et al., 2013; Sesa-Ashton et al., 2022). These findings suggest a functional loop wherein cortical regions modulate sympathetic outflow, and peripheral autonomic signals, in turn, influence central processing.

Specifically, during heartbeats, spontaneous bursts of muscle sympathetic nerve activity (MSNA) – which regulate arteriolar diameter in skeletal muscles and contribute to beat-to-beat blood pressure control – exhibit strong temporal coupling within the dlPFC and MCC activation, as measured using concurrent microneurography and fMRI (James et al., 2013). This interaction suggests a potential top-down modulation of sympathetic output to skeletal muscle vasculature. Furthermore, transcranial alternating current stimulation of the dlPFC in awake humans has been shown to partially entrain MSNA, heart rate, and blood pressure, underscoring the dlPFC’s role in autonomic regulation and cardiovascular control (Sesa-Ashton et al., 2022).

Complementing these findings, research by Riaz et al. has demonstrated a correlation between MSNA inhibition and beta-band rebound activity in the ACC and somatosensory cortex (Riaz et al., 2022). This suggests that sensorimotor processing contributes to the modulation of sympathetic responses, possibly through the integration of afferent somatosensory input with autonomic regulation.

The observed synchronization in response to MSNA suppression supports the notion that beta suppression may be related to the brain’s engagement in processing sensory input about cardiac sensations, which are generated by a cascade of peripheral reactions. This engagement in afferent information processing may be particularly pronounced during interoceptive attention – a cognitive state known to modulate cortical oscillations, particularly in the beta frequency band, thereby enhancing the integration of internal bodily signals with autonomic regulation (Angioletti & Balconi, 2022).

The convergence of these results strengthens the idea that oscillatory responses tied to the cardiac cycle carry behavioral significance. In our study, the fact that the beta suppression tracks interoceptive accuracy implies that this neural measure indexes the efficacy of interoceptive signal processing. Our findings suggests beta suppression reflects individual variability in interoceptive processing, potentially serving as a candidate neural marker for further investigation. Those with a more pronounced suppression appear to have more precise or sensitive internal monitoring, resulting in higher interoceptive performance. This insight has potential implications for using neural oscillatory markers to assess or even train interoceptive abilities in the future.

## Materials and methods

### Participants

51 volunteers participated in the experiment (21 men and 30 women), aged 33± 6.7 years (M ± SD) with 19 – 62 range. Exclusion criteria included a history of cardiovascular or respiratory disease, head trauma, intellectual disability, neurological or psychiatric condition, use of medication affecting the nervous system, and claustrophobia. Additionally, participants included in this study had normal or corrected-to-normal hearing and vision.

Ethical approval was received from the local Ethics Committee of the University of Jyväskylä and all the study procedures were conducted in accordance with the Declaration of Helsinki. Prior to any study procedures, participants were informed about the study, and a written informed consent was obtained from all participants.

### Procedure

Participants performed modified heartbeat and an auditory discrimination task (HDT) (Critchley et al., 2004a; Lyyra & Parviainen, 2018). The task included randomized trials that required participants to focus attention on their own heartbeat (interoceptive attention task) or a sound stimulus (exteroceptive attention task). Both task conditions were based on monitoring a sequence of sounds that were played either simultaneously or with a delay with respect to individual’s own heartbeat. Additionally, in all the task trials the pitch of the sound was constant (800 Hz, 100 ms).

During each trial, a visual cue was displayed on the screen to indicate the task condition. A heart icon signaled the interoceptive attention task, instructing participants to focus on whether in given trial the sounds were played simultaneously with their heartbeat. After the sound sequence ended (20 heart beats), they responded by pressing the yellow button if the sound was subjectively felt to be in sync with their heartbeat and the blue button if it was out of sync (asynchronous).

A musical note icon indicated the exteroceptive attention task, requiring participants to focus on the pitch of the sound. After the sequence ended, they pressed the yellow button if they detected a pitch deviation and the blue button if the pitch remained unchanged (Fig. 1A). Importantly, auditory sounds were present during both types of attention blocks to prevent any differences related to sound response between tasks. The auditory stimulus was triggered by the onset of the participant’s heartbeat either simultaneously (synchronized condition: 200 ms delay after R-peak), or with a delay (asynchronized condition: 500 ms delay after R-peak). The 200 ms delay for the simultaneous condition was selected based on subjective assessments, as it corresponded to the highest synchronicity judgments, while the 500 ms delay corresponded to the lowest (Garfinkel et al., 2015; Wiens & Palmer, 2001).This delay also corresponds to the time interval between the R-wave peak and the base of the leading edge of the finger pulse wave (Payne et al., 2006). In both tasks, participants completed 6 blocks each containing 20 heartbeats.

In this study, the MEG data analysis was based on the interoceptive and exteroceptive tasks with simultaneous sound presentation to avoid overlap between induced activity and phase-locked activity related to the auditory response. Asynchronous and synchronous conditions in the interoceptive task were used to assess interoceptive accuracy.

### ECG data recording

ECG was measured using Ag/AgCl electrodes placed at the fifth left intercostal space at the midclavicular line and the right sternum (with the ground electrode on the participant’s right front side below the ribs). The position of the electrode on the sternum was adjusted to ensure a clear R-peak of the QRS complex was detectable in the ECG signal, which was acquired using a specially designed device controlled by ARDUINO® software. Custom software script detected individual R-peaks and controlled the presentation of heart-beat locked auditory stimuli. The device also sent a trigger marking each heartbeat to the trigger channel in MEG data. The resulting triggers associated with the R-peak were then applied to define epochs for further signal analysis.

### MEG data recording

MEG data was collected using a whole-head 306-sensor (102 magnetometer channels and 204 planar gradiometer channels) Elekta Neuromag TRIUX system (MEGIN Oy, Helsinki, Finland) in a magnetically shielded sound-attenuated room. Data were sampled at 1000 Hz and filtered at 0.1-330 Hz. We measured participants’ head shapes using five head position indicator (HPI) coils that were attached to the scalp (three HPI coils placed on the forehead and one behind each ear).

### MEG data preprocessing and epoch selection

The MEG signal was preprocessed using MaxFilter software (v.2.2) to reduce external noise through the temporal signal-space separation (tSSS) method (Taulu et al., 2005). Additionally, head movements were compensated by realigning the head position at each time point to an “optimal” common reference position (head origin).

For further offline analysis, we utilized the MNE-Python (v 1.7) software (Gramfort et al., 2013). The raw data contaminated with myogenic artifacts were automatically detected using MNE-python function ‘annotate_muscle_zscore’ with default parameters (ch_type =‘mag’, threshold = 5.0, min_length_good = 0.2, filter_freq = [110, 140]) and excluded from analysis. After its raw data were band-passed in 1 - 40 Hz range and resampled offline to 300 readings per second.

The biological artifacts were removed from the continuous data using the fastICA method implemented in MNE-Python (Hyvärinen, 1999). Because temporal signal space separation induces rank deficiency, we defined the maximum number of ICA components as the rank of raw data. We used a semi-automated approach implemented in the MNE-python for detection and removal of electrocardiographic (ECG) and vertical electrooculographic (EOG) artefacts. To ensure that the automatically detected ICA components were caused by the artefacts, we checked their timecourses and spatial distributions. The number of cardiac components removed from the data was 2.3+0.2. We used only gradiometers for analysis because they are less sensitive to fluctuations in stroke volume and cardiac field artefacts (Buot et al., 2021).

We extracted R peak locked epochs from −0.500 to 1.500 s relative to the R peaks of the heart beats. Epochs with a cardiac cycle shorter than 0.7 s were excluded from analysis.

### Time-frequency analysis of the MEG data at the source space

Source reconstruction was performed in a template brain (FreeSurfer fsaverage-5.1.0 template) (Fischl, 2012). Coregistration between the T1-weighted MRI template scalp and the digitized scalp was performed for each participant using a three-axis scaling mode.

To compute the β-band (16–28 Hz) power we used ‘mne.time_frequency.tfr_morlet’ function (implemented in the MNE-Python open-source software), with Morlet wavelets using adaptive Morlet constant for each frequency bin (frequency/2) and 2 Hz steps.

A source reconstruction was performed using ‘mne.minimum_norm.apply_inverse_tfr_epochs’ function, and standardized low-resolution brain electromagnetic tomography (sLORETA) localization method with a source space consisting of 4,098 vertices per hemisphere (Pascual-Marqui, 2002).

The empty room recording was used to construct the noise covariance matrix. The empty room MEG recordings were spatially filtered using the tSSS method and signals were passband 1–40 Hz.

The data were baseline-corrected by dividing β-power at each time point by the averaged power computed over the pre-stimulus interval (from −0.300 to −0.100 s relative to R-peak onset). The resulting values were converted to dB by log10-transformation and multiplication by 10 (Keil et al., 2022).

### Statistical analysis. Regions of interest and time interval selection

The selection of regions of interest (ROIs) we employed a hybrid approach initially identifying anatomical ROIs based on existing literature and subsequently conducting statistical analyses to verify the presence of significant changes in these regions.

It is known that regions such as the insula, anterior and mid-cingulate cortex, medial prefrontal areas, and somatosensory and motor cortex are involved in processing interoceptive signals (Craig, 2009; Critchley & Harrison, 2013; Khalsa et al., 2009; Uddin, 2015). Therefore, we selected these regions as potential areas involved in cardiac interoception.

We then tested whether power in these ROIs significantly differed from baseline. This was done using a fully flattened average approach applied to data from both the exteroception and interoception tasks under the simultaneous condition (Bowman et al., 2020). To rule out the possibility of missing areas with significant R-peak-related power changes, we combined the preselected ROI approach with a data-driven approach. Therefore, we performed spatiotemporal analysis on all vertices to identify the time and source distribution of the peak activity. We applied one-sample permutation test with spatiotemporal threshold-free cluster enhancement (TFCE) across time (from 0.100 to 1.0 s) and space (8196 vertices) using mne.stats.permutation_cluster_1samp_test (n_permutations=1000) and starting threshold h0 = 0, step size dh = 0. 4. To minimize computational costs, we applied a stringent cluster-forming threshold based on the t-distribution with a p-value of 0.05 for data with spatial adjacency.

Next, to precisely refine the boundaries of the preselected ROIs, we overlaid the obtained activity distribution in the significant TFCE cluster with functional anatomical regions from the multi-modal atlas from the Human Connectome Project HCPMMP1_combined and retained only statistically significant vertices (Glasser et al., 2016). To compare the activity between the interoceptive and exteroceptive tasks, a paired t-test was used with effect size estimation using Cohen’s d. For comparing activity at rest and during the performance of exteroceptive and interoceptive tasks, repeated measures ANOVA (rmANOVA) was used with partial eta squared (ηp2) for effect size estimation. Pairwise comparisons were conducted using Holm–Bonferroni post-hoc tests.

### Interoception accuracy

During the experiment, as part of the interoceptive task, participants were required to determine whether the auditory tones were played simultaneously with their heartbeat or not after each block of 20 consecutive tones. Interoceptive accuracy was assessed as the ratio of correct responses to the total number of interoceptive blocks. To investigate the relationship between interoceptive accuracy and MEG signal power, Spearman’s correlation was used.

## Acknowledgments

We thank Doris Hernández Barros for her assistance with data collection.

## Author Note

We have no known conflict of interest to disclose.

