## Supporting information for "Oscillatory markers of interoceptive attention: beta suppression as a neural signature of heartbeat processing"

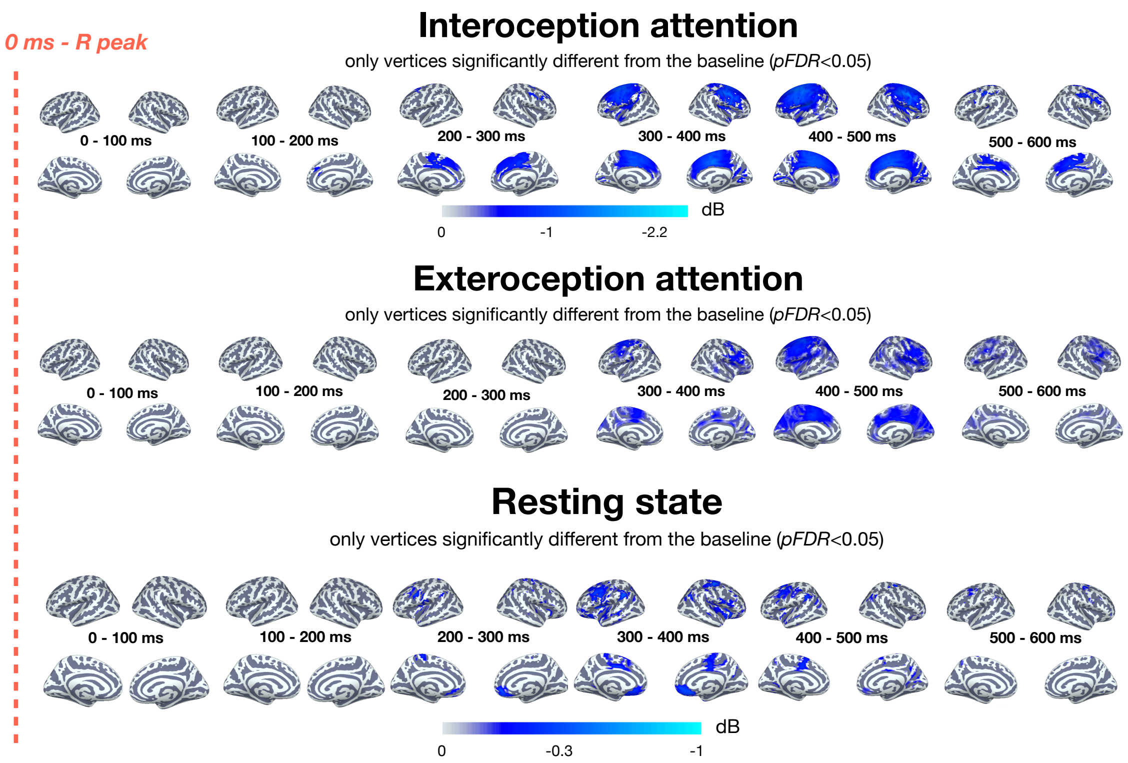


**S1 Fig.** Source-level distribution of beta power dynamics during resting state, interoceptive, and exteroceptive conditions, computed in 100 ms. Displayed vertices reflect significant deviations from baseline ($p < 0.05$, one-sample t-test, FDR corrected).

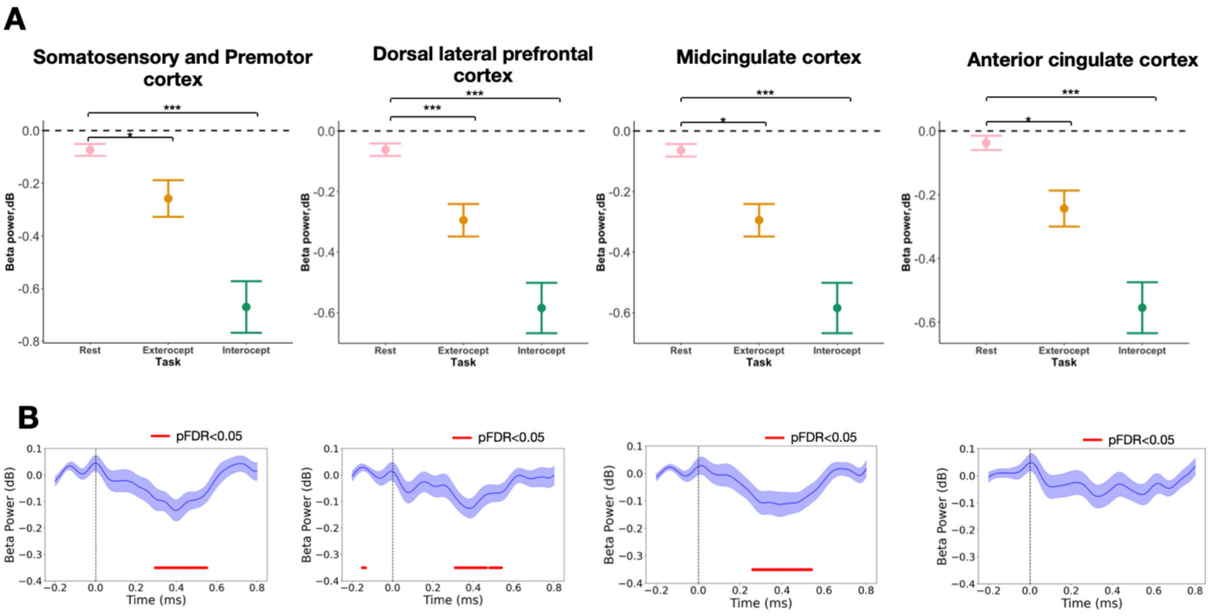


**S2 Fig.** Beta power during resting state in ROIs. (A) Beta power in resting state and during the HDT conditions in ROI averaged in 0.310 to 0.530 s after R-peak (Points and error bars on graphs represent M ± SEM in all subjects. * $p < 0.05$, ** $p < 0.01$, *** $p < 0.001$ (rmANOVA, Holm correction test)); (B) Time course of β power changes during the resting state and ROIs.

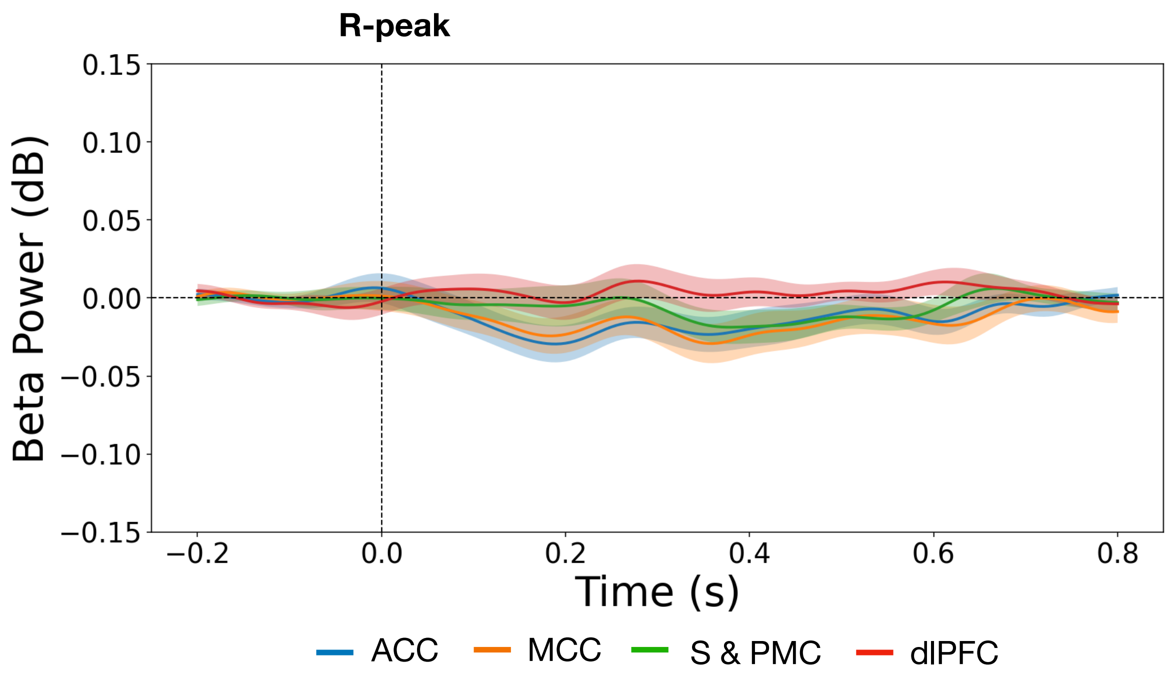


**S3 Fig.** R-peak-related beta power change in ROIs in surrogate R-locked epochs

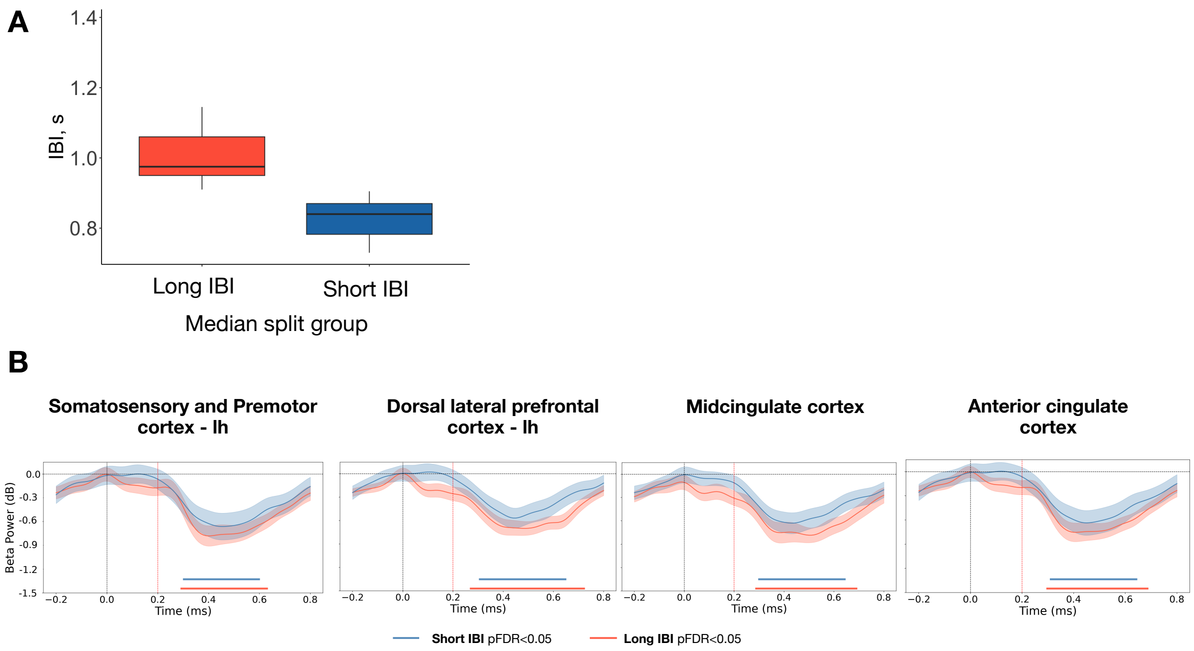


**S4 Fig.** R-peak-related beta power change for all tasks collapsed in participants with long and short heart rates.
